# A lipid signature of BAK-driven apoptotic pore formation

**DOI:** 10.1101/2024.10.16.618570

**Authors:** Rachel T. Uren, Matthew E. Ritchie, Agnes W. Wong, Justin P. Ludeman, Etsuko Uno, Vinod K. Narayana, David P. De Souza, Dmitri Sviridov, Ruth M. Kluck

## Abstract

Apoptotic cell death is regulated by the BCL-2 protein family, with clusters of BAK or BAX homodimers driving pore formation in the mitochondrial outer membrane via a poorly understood process. There is growing evidence that, in addition to BAK and BAX, lipids play an important role in pore formation. Towards a better understanding of the lipidic drivers of apoptotic pore formation in isolated mitochondria, two complementary approaches were taken. Firstly, the lipids released during BAK-mediated pore formation were measured with targeted lipidomics, revealing enrichment of long chain polyunsaturated lysophospholipids (LPLs) in the released fraction. In contrast, the BAK protein was not released suggesting that BAK and LPLs locate to distinct microdomains. Secondly, added cholesterol not only prevented pore formation but prevented the clustering of BAK homodimers. Our data lead us to a model in which BAK clustering triggers formation of a separate microdomain rich in LPLs that can progress to lipid shedding and the opening of a lipid-lined pore. Pore stabilisation and growth may be due to BAK dimers then moving to the pore edge. Our BAK-lipid microdomain model supports the heterogeneity of BAK assemblies, and the observed lipid-release signature gives new insight into the genesis of the apoptotic pore.

## INTRODUCTION

A key decision point during apoptotic cell death is the rupture of the mitochondrial outer membrane (MOM) triggered by effector proteins of the BCL-2 family, BAK (BCL-2 homologous antagonist/killer) and BAX (BCL-2-associated protein X) ^1, 2^, and perhaps BOK (BCL-2-related ovarian killer) ^3^. Mitochondrial outer membrane permeabilization (MOMP) releases cytochrome *c* and other proteins from the mitochondrial intermembrane space into the cytoplasm, leading to the activation of caspases and loss of cell viability. Here we refer to the MOM rupture that is dependent on BAK or BAX as the “apoptotic pore”. Importantly, the molecular mechanism of apoptotic pore formation remains unclear (reviewed in ^4^).

BAK and BAX each comprise nine α-helices, with marked structural and sequence homology ^5, 6^. Initially, non-activated BAK is a globular protein anchored to the MOM by its C-terminal transmembrane domain ^7^, whereas BAX is largely cytosolic until it translocates to the MOM ^8-10^. Upon an apoptotic trigger, initiators of the BCL-2 family (e.g. BID, BIM) are upregulated and then transiently bind to BAK or BAX to trigger a series of conformational changes including dissociation of the N-and C-termini to expose the BAK or BAX BH3 domain. The exposed BH3 domain of activated BAK or BAX can then engage the hydrophobic binding groove of another activated monomer on the membrane to create a closed stable symmetric homodimer ^11, 12^.

Structural studies of truncated BAK and BAX show that α2-α5 core dimers are rigid and present a hydrophobic surface on one face ^13-17^. The BAK α2-α5 core dimers also showed binding sites for phospholipid head groups and acyl chains, with lipid “bridging” between dimers ^15^. Related lipid binding sites were not evident in BAX α2-α5 core dimers although association with lipids was inferred as BAX structures were dependent on the lipid environment ^17^. In simulations the BAK α2-α5 core dimer perturbs lipid bilayers causing increased membrane curvature and bilayer thinning ^15^, together with aggregation of triacylglycerides (TAGs) under the BAK α2-α5 core dimer and compensatory changes in the lipid distribution in the surrounding membrane ^18^.

Mitochondrial studies show that beyond the α2-α5 core regions, the N- and C-termini in BAK and BAX homodimers are relatively flexible ^7, 11-14, 19-22^. While the N-terminus makes no contact with protein or membrane, the C-terminal amphipathic helices α6, 7 and 8 lie in-plane with the membrane suggesting that this region may also perturb the membrane bilayer ^21^.

Membrane-embedded BAK dimers adopt disordered clusters on the MOM, as various contact points between dimers have been captured with cysteine linkage or magnetic distance measures but none appear essential for clustering or pore formation ^19, 23, 24^. Consistent with multiple contact points, BAK and BAX dimers coalesce into various architectures including clusters, lines, rings or arcs (reviewed in ^25^). In addition, if cells express both BAK and BAX, the proteins form mixed unordered assemblies of variable dimensions that completely or partially encircle the pore ^26^. Collectively, these data argue that the apoptotic pore is not a rigid proteinaceous structure but contains both protein and lipid elements ^27, 28^.

Consistent with the apoptotic pore being at least partly lipidic is the ability of the MOM phospholipid composition to influence apoptosis. Membrane composition can alter several steps in BAK and BAX function, including their translocation to mitochondria, activation, dimerisation and pore formation ^27, 29-37^. Membrane changes may also be induced by apoptotic signalling, as lysophospholipids accumulated in mitochondria during Fas signalling ^38^, modelling showed phospholipid distribution changed in response to BAK homodimers ^18^, and unsaturated phospholipids were enriched in BAK oligomers extracted from mitochondria ^39^.

Directly measuring MOM lipid changes during pore formation is challenging due to the small size and complex membrane architecture of mitochondria. The MOM is difficult to separate from the MIM, and the MOM lipid composition is distinct from that of other cellular compartments and even distinct from the inner mitochondrial membrane ^40-45^.

Here we examined whole isolated mitochondria to further understand the involvement of lipids in apoptotic pore formation in a native context. We employed targeted lipidomics to quantify lipids released from isolated mitochondria during pore formation, reasoning that released material may reflect a lipid defect from which the pore emerges. Moreover, studying isolated mitochondria allowed examination of lipid redistribution events in the absence of significant lipid metabolism or lipid transfer between organelles. Notably, BAK-driven pore formation was associated with release of lysophospholipids but not BAK, suggesting the presence of discrete microdomains of lysophospholipids and of BAK, and a pore opening closest to the site of lysophospholipid aggregation rather than BAK clustering. Exogenous cholesterol was able to prevent BAK clustering and pore formation, indicating the importance of microdomains in apoptotic pore formation.

## RESULTS

### Apoptotic pore formation in isolated mouse liver mitochondria

To explore the interplay between BAK and membrane lipids, mouse liver mitochondria (MLM) were freshly isolated ^46^. In this system, fractionation removes cytosolic BAX, allowing BAK function to be studied in the absence of BAX. As previously ^46^, incubation with caspase-cleaved BID (cBID) released essentially all cytochrome *c* into the supernatant (Fig. 1a) and activated all BAK (Fig. 1b), which then assembled into higher order complexes (Fig. 1c). Time course experiments demonstrated significant BAK activation at 2 min (Fig. 1b), high order BAK clusters at 15 min (Fig. 1c) and full cytochrome *c* release at 20 min (Fig. 1a).

**Figure 1.**
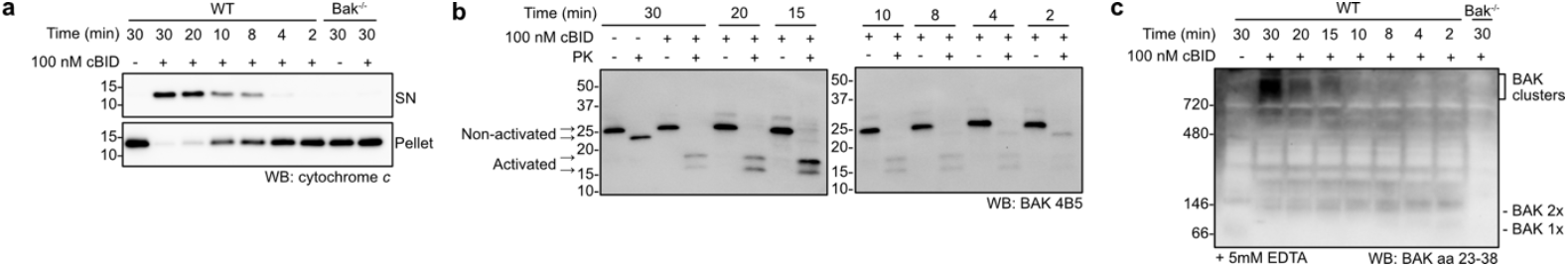
Apoptotic pore formation in mitochondria as shown by cytochrome c release, BAK activation and BAK clustering. WT or Bak^-/-^ MLM were incubated with cBID at 37°C and sampled at the indicated times. **(a)** Robust cytochrome *c* release is dependent on the presence of both cBID and BAK. Pellet and supernatant (SN) samples were immunoblotted for cytochrome *c*. **(b)** BAK activation is triggered by cBID as shown by increased susceptibility to proteinase K digestion. Aliquots removed prior to fractionation were treated with proteinase K and immunoblotted with antibody to the BAK BH3 domain (clone 4B5). After proteinase K, non-activated BAK runs as a ∼23 kD fragment, while activated BAK runs as ∼17 kD and ∼15 kD fragments. **(c)** High molecular weight BAK complexes (“BAK clusters”) form in the presence of both cBID and BAK. The pelleted mitochondrial fractions from (a) were run on Blue Native PAGE and immunoblotted for BAK (aa23-38). Addition of 5 mM EDTA immediately prior to BNP slightly enhances the detection of mouse BAK under BNP conditions, providing some evidence of dimers (2X species) forming after cBID, but decreasing as high molecular weight complexes form (see lane 2, 30 min incubation). Data are representative of three independent experiments.

### BAK is retained in mitochondria following pore formation

To examine the release of lipids as well as proteins we employed a two-step centrifugation protocol (Fig. 2a). After pore formation initiated by cBID, MLM were centrifuged at 10,000 *g*, and the resulting supernatant (SN^10^) subjected to 100,000 *g*, generating a secondary pellet fraction (Pellet^100^). The Pellet^100^ fraction was the optimal fraction for lipidomics analysis as it removed buffer constituents (such as HEPES) that interfere with mass spectrometry and also concentrated the sample to improve detection (see Discussion).

**Figure 2.**
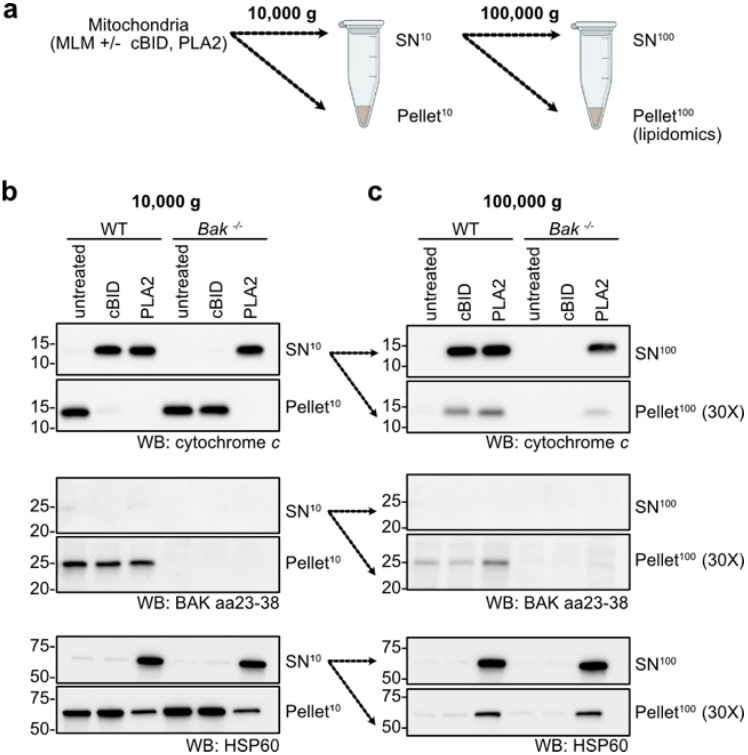
BAK is retained in the mitochondrial outer membrane during pore formation. **(a)** Two-step centrifugation used to fractionate proteins and lipids released from mitochondria during apoptotic pore formation. WT or *Bak*^*-/-*^ MLM were incubated with cBID (100 nM) or PLA2 (5 μg/mL) at 37°C for 30 min. The indicated fractions were collected for western blot and lipidomics. **(b)** 10,000 *g* fractions show that pore formation triggered by cBID released cytochrome *c* but not BAK or HSP60. **(c)** 100,000 *g* fractions show only background levels of BAK in Pellet^100^ after cBID treatment (compare lanes 1 and 2). Note that Pellet^100^ samples are ∼30X more concentrated than SN^100^ samples. Data are representative of three independent experiments.

As a negative control for pore formation, MLM from *Bak*^-/-^ mice ^2^ were tested in parallel with MLM from wildtype mice. As a positive control for liberation of lipids independent of any apoptotic signalling, MLM from both genotypes were separately incubated with the phospholipase A2 (PLA2) from bee venom ^47^ to digest the mitochondrial membrane.

While cBID failed to release cytochrome *c* from *Bak*^-/-^ MLM, PLA2 caused cytochrome *c* release from both genotypes (Fig. 2b), indicating that the lipase was acting in a BAK-independent manner. PLA2 also partially digested the inner membrane, as shown by the release of the matrix protein HSP60 (Fig. 2b). Notably, whereas a portion of BAK was liberated by PLA2 treatment, BAK was not released during apoptotic pore formation (see Pellet^100^, Fig 2c). Thus, BAK is membrane-integrated in healthy mitochondria, and remains integrated and associated with the mitochondrial pellet upon apoptotic pore formation.

As bee venom PLA2 is a calcium-dependent enzyme, the calcium chelator EDTA prevented PLA2-mediated release of cytochrome *c* and HSP60 (Fig. 3). In contrast, EDTA had no effect on BAK-mediated cytochrome *c* release (Fig. 3), indicating that the apoptotic pore forms independently of calcium-dependent enzymes like PLA2.

**Figure 3.**
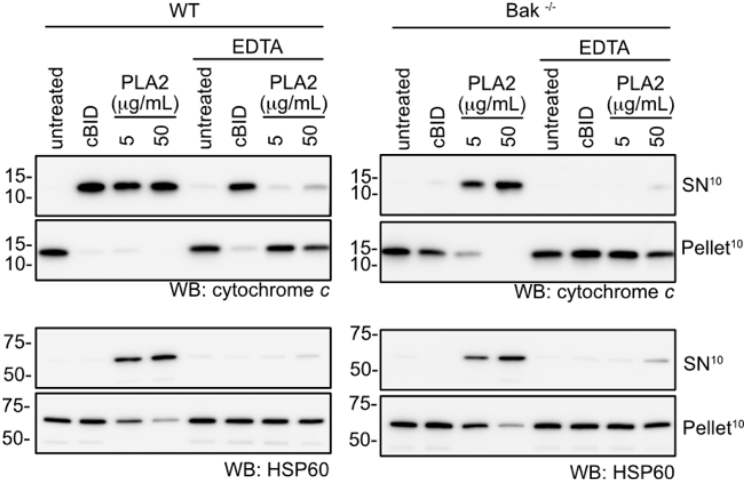
BAK-mediated pore formation is calcium-independent. Prior to incubation of WT or Bak^-/-^ MLM with cBID (100 nM) or PLA2 (5 or 50 μg/mL), MLM were incubated with EDTA (20 mM) to sequester residual calcium in the reaction buffer, and thus inhibit any calcium-dependent enzymes such as PLA2. Pellet^10^ and SN^10^ fractions were western blotted for cytochrome *c* or HSP60. Data are representative of three independent experiments.

### Targeted lipidomics to identify lipids liberated from apoptotic mitochondria

To test for release of lipids from mitochondria, Pellet^100^ fractions were collected from six repeat experiments, each experiment comparing liver mitochondria from one WT and one *Bak* ^-/-^ mouse under three different treatments (untreated, cBID, or PLA2) (Table S1). The expected cytochrome *c* release was confirmed by western blot analysis of the Pellet^10^ and SN^10^ fractions (Fig. S1). The Pellet^100^ samples were then analysed by mass spectrometry using a targeted lipidomics panel of 495 lipid metabolites including 4 internal standards. A total of 348 (70%) metabolites were detected (Table S2; MassHunter, Agilent), demonstrating excellent coverage of this lipid panel. Further lipidomic analysis including interactive summary plots can be found at https://github.com/mritchie/BAKLipidomicAnalysis/.

According to the raw lipid counts the majority of lipids in the Pellet^100^ fractions from WT Untreated MLM were PC or PE species (Table S3 and S4). PC and PE are the major components of all cellular membranes including the MOM ^42, 45^, indicating recovery of membrane fragments of some kind in the Pellet^100^ fractions. The very low abundance of cholesterol (<0.01 %) is consistent with a mitochondrial origin, as cholesterol levels in the mitochondria are low relative to other cellular membranes ^42, 45, 48^.

To summarise changes in lipid release across all samples, a multidimensional scaling (MDS) analysis was performed. A distinct cluster was obtained from PLA2 samples (Fig. S2). PLA2 cleaves di-acyl phospholipids (e.g. PC and PE) at the sn-2 acyl linkage to generate free fatty acids and lysophospholipids (Fig. S3a and reviewed ^49-51^). Accordingly, fold-change analysis revealed lysophospholipids (e.g. LPC and LPE) were increased in the Pellet^100^ fractions of PLA2 treatments, denoting a signature of released lysophospholipids (Fig. S4). A corresponding decrease of substrate phospholipids (such as PC and PE) in the Pellet^100^ fractions was also evident (Fig. S4). Parallel production of free fatty acids by PLA2 is assumed to occur, but they are not measured by the lipidomics assay. Together, these findings indicate that PLA2 successfully hydrolysed a range of phospholipids in MLM, with expected lysophospholipid products released from MLM and recovered in the Pellet^100^ fractions.

### Specific release of long chain polyunsaturated lysophospholipids during pore formation

To test for lipids released after BAK activation and pore formation, we next compared the lipidomic profiles of the Pellet^100^ samples from WT MLM treated with cBID versus WT Untreated controls. The “top ten” over-represented or under-represented lipids were determined by ranking data by raw *p*-value, and then classifying by log2fold-change as over-represented lipids (positive log2fold-change) or under-represented lipids (negative log2fold-change) (Fig. 4, Tables 1 and 2). A full list of fold-changes across all 348 of the detected lipids, for all contrasts, is available in Table S6. *P*-values were adjusted for multiple testing using the false discovery rate (FDR) method, and those with a FDR < 0.05 were considered differentially abundant. There was one significantly altered lipid, lysophosphatidylcholine (LPC 22:5(104)), FDR = 0.03224) that was significantly increased in Pellet^100^ fractions from cBID treated WT MLM compared to Untreated controls (Fig. 4a, Table 1 and Table S6).

**Table 1:**
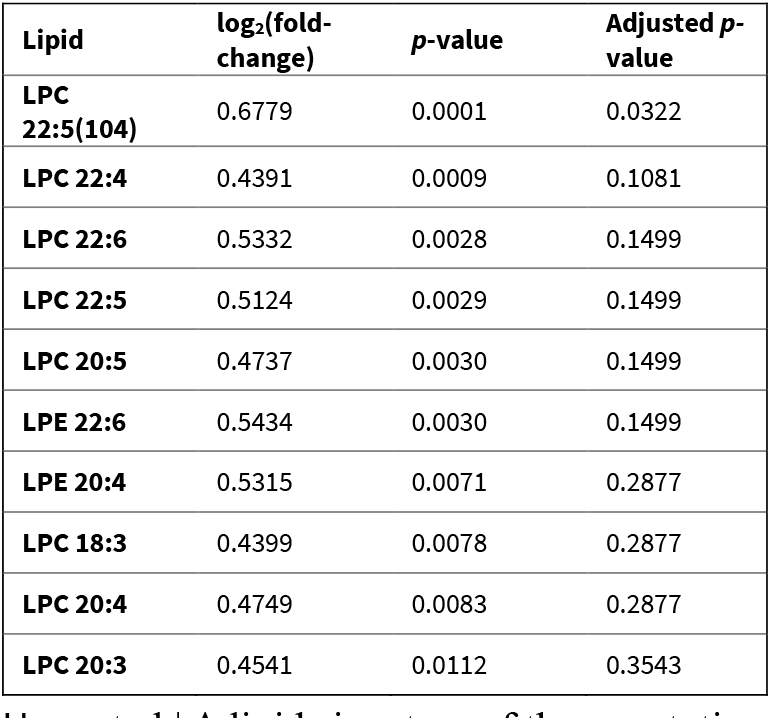
Over-represented lipids WT cBID versus WT Untreated (ranked by *p*-value)

**Table 2:** Under-represented lipids WT cBID versus WT Untreated (ranked by *p*-value)

| Lipid | log<sub>2</sub>(fold-change) | *p*-value | Adjusted *p*-value |
| --- | --- | --- | --- |
| **TG 18:0_18:0_18:0** | -0.5944 | 0.0009 | 0.1081 |
| **Cer(d18:1_26:0)** | -0.3940 | 0.0223 | 0.4755 |
| **dhCer 24:1** | -0.2280 | 0.0547 | 0.9068 |
| **Cer(d17:1_24:0)** | -0.2568 | 0.0729 | 0.9957 |
| **PE(P-16:0_18:1)** | -0.1433 | 0.1104 | 0.9957 |
| **Cer(d16:1_24:0)** | -0.2529 | 0.1136 | 0.9957 |
| **Hex1Cer (d18:1_24:0)** | -0.2091 | 0.1223 | 0.9957 |
| **PC 34:0** | -0.1547 | 0.1792 | 0.9957 |
| **SM 44:1** | -0.1146 | 0.1961 | 0.9957 |
| **SM 41:1** | -0.1272 | 0.2160 | 0.9957 |

**Figure 4.**
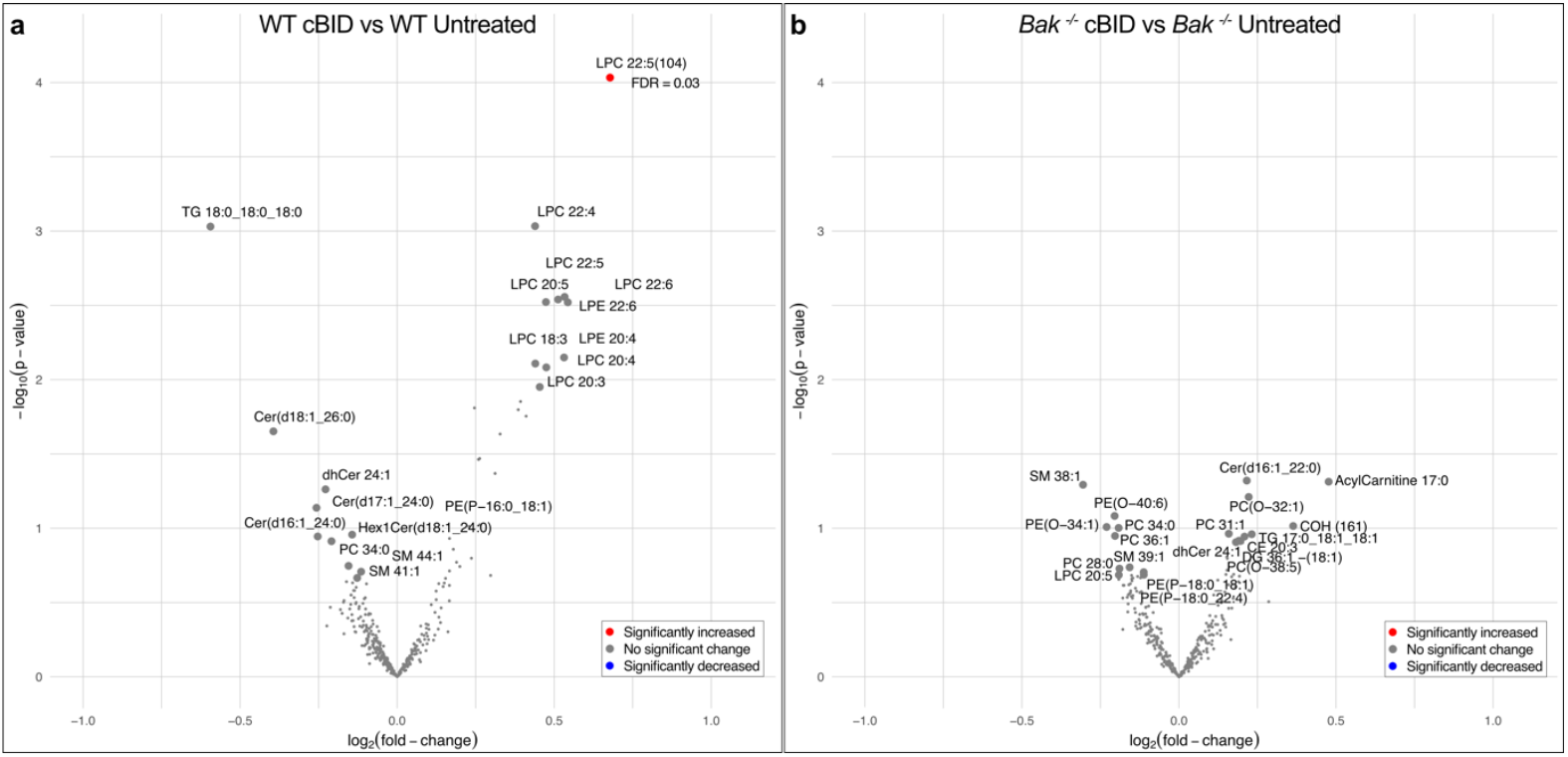
Specific release of long chain polyunsaturated lysophospholipids (LPLs) from mitochondria during pore formation. Volcano plots compare Pellet^100^ fractions from **(a)** WT cBID versus WT Untreated samples and **(b)** *Bak*^-/-^ cBID versus *Bak*^-/-^ Untreated samples. Lipids with FDR <0.05 are highlighted in red (increased, i.e. over-represented) or blue (decreased, i.e. under-represented). Lipid log_2_(fold-change)(x-axis) is plotted against (-log_10_(p-value)(y-axis). The 10 most over-represented or under-represented lipids (as ranked by *p*-value) are labelled and highlighted with enlarged data points. (See Table S6 for all contrasts, and Tables 1 and 2 for 10 most over-represented or under-represented lipids in the WT cBID versus WT Untreated contrast).

Remarkably, related lysophospholipids accounted for all ten of the most increased lipids liberated from apoptotic mitochondria as all were long chain (>20 carbons) and polyunsaturated (3-5 double bonds) (including LPC 22:5, an isomer of the top-ranked LPC 22:5(104)). This polyunsaturated lysophospholipid signature was not observed when comparing *Bak*^-/-^ cBID treated samples with *Bak*^-/-^ Untreated controls (Fig. 4b, Table S6), further supporting the specificity of this signature for BAK-dependent pore formation. An important observation is that the lysophospholipids liberated with apoptosis were longer and more unsaturated than those liberated by PLA2 treatment (Fig. S4, Table S6), again indicating distinct modes of lipid release by the two treatments (see Discussion).

To gain a wider view of the behaviour of major lipid classes during apoptotic pore formation, lipid set enrichment analysis of the lipidomics output was performed by two different approaches: a specific analysis classified by lipid headgroup (ROAST ^52^)(Fig. 5a) or by a comprehensive analysis covering the biophysical, chemical and cell biological features of each lipid molecule (LION/web ^53, 54^) (Fig. 5b). ROAST analysis showed enrichment of lysophospholipids with a phosphatidylcholine headgroup (LPC lipid class) following cBID treatment in the increased category (*p*-value = 0.029) (Fig. 5a), consistent with eight of the top ten over-represented lipids being LPC lipids (Table 1). After PLA2 treatment there was enrichment for LPC, LPE and LPI headgroup classes amongst increased lipids (Fig. 5a), again highlighting the distinct lipid signatures of apoptotic pore formation and PLA2 digestion (see Discussion). The LION/web analysis confirmed enrichment of lysoglycerophospholipids (which includes LPC) (Fig. 5b). In addition, lipids in the category of “positive intrinsic curvature” were enriched, consistent with LPC promoting positive curvature ^55^. Positive intrinsic curvature is a key feature of lipids that can promote membrane budding and vesicle release.

**Figure 5.**
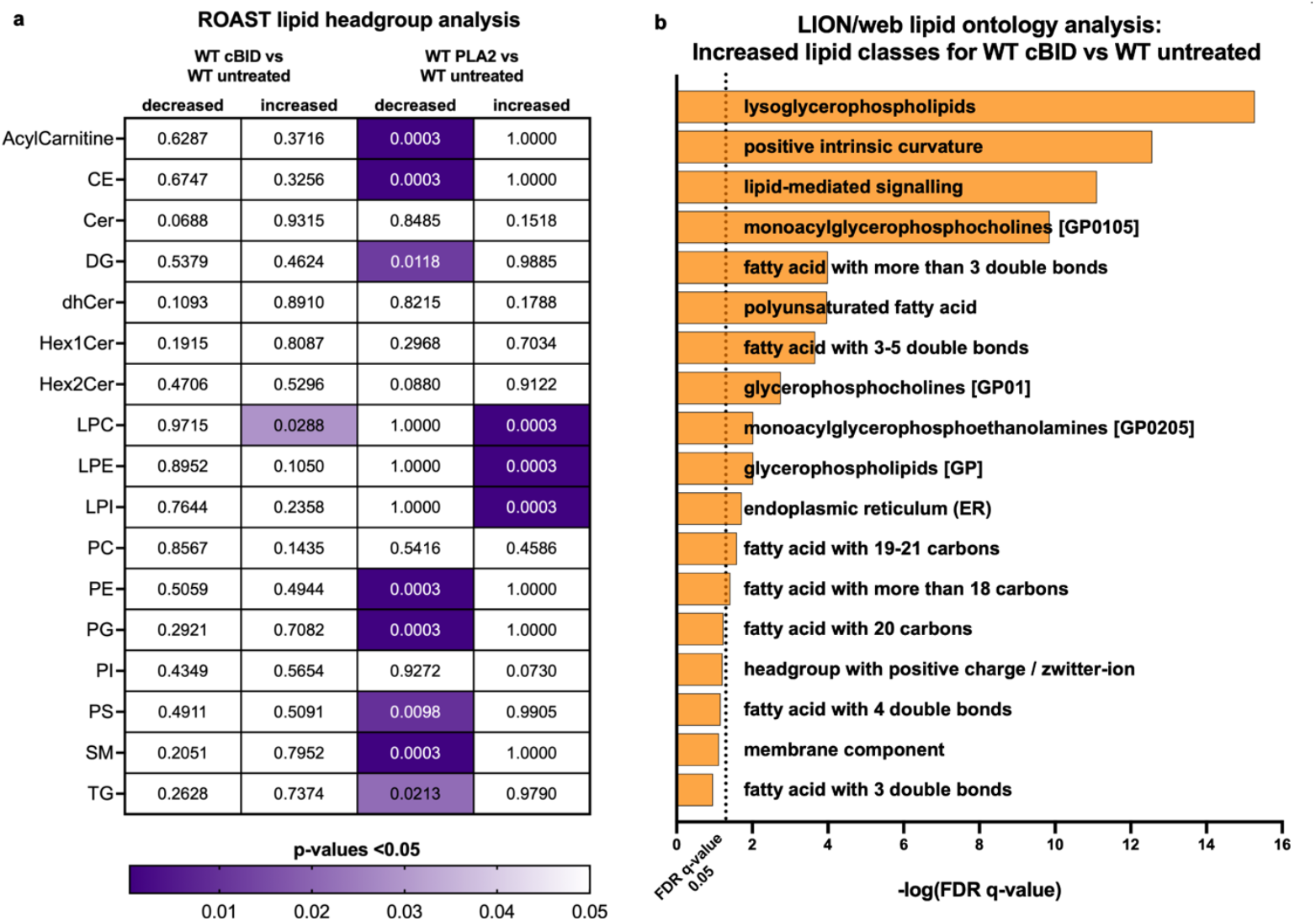
Enrichment analysis of lipid classes detected with apoptotic signalling. **(a) Preference for phosphatidylcholine headgroup in lysophospholipids released during apoptotic pore formation.** An enrichment analysis based on the major lipid headgroup classes using ROAST (see Methods) was performed for Pellet^100^ fractions with contrasts indicated (WT cBID vs WT Untreated or WT PLA2 vs WT Untreated) (see also Table S7 for all contrasts). ROAST enrichment *p*-values are shown for each lipid headgroup testing for decreased or increased lipid classes. Significant *p*-values (<0.05) are highlighted (purple gradient). Also note the distinct signatures for cBID compared to PLA2. **(b) Preference for lipids with positive intrinsic curvature in lipids released during apoptotic pore formation**. A complementary enrichment analysis encompassing a range of lipid features using the LION/web lipid ontology tool (see methods) was performed. The top 26 lipids (raw *p*-value < 0.2 (Supp. Table 8)) with positive log_2_fold-change from the WT cBID vs WT Untreated contrast were input as the ‘lipid target list’ and the full list of detected lipids (384) were input as the ‘lipid background list’. “Lysoglycerophospholipids” (which encompasses LPC), “lipid-mediated signalling” and “positive intrinsic curvature” were the top 3 category descriptors. A vertical dotted line indicates the FDR q-value significance cut off of 0.05.

In summary, lipidomics identified for the first time a lipid signature of long chain polyunsaturated lysophospholipids (LPLs) specifically released during apoptotic pore formation. The specificity of the lipids being released argues that bulk membrane changes such as lateral lipid redistribution into microdomains (e.g. enriched for lysophosholipids) is occurring during pore formation, and possibly driving one or more steps leading to pore formation. As lysophospholipids are known to position at the edge of lipidic pores, our data support previous evidence that apoptotic pores are lipidic in nature. As BAK is not co-released with the lysophospholipids, BAK does not directly bind to them and may not be present in the microdomain from which these lipids are released.

### Triacylglycerides are retained in mitochondria following apoptotic pore formation

While lysophospholipids were relatively over-represented in the Pellet^100^ fraction following pore formation, other lipids were relatively under-represented (Fig. 4a, Table 2, Table S6), likely due to enhanced retention in the MOM upon pore formation. The ten lipid species most under-represented in Pellet^100^ in the cBID/BAK group included triacylglycerides and ceramides (Table 2). The most under-represented was the triacylglyceride TG 18:0_18:0_18:0 which reached a log2fold-change of -0.59, (FDR = 0.11, ns). These lipids may be retained in the MOM by being attracted to other domains. Indeed, triacylglycerides were seen to focus beneath BAK α2-α5 core dimers in molecular dynamics simulations ^18^ (see Discussion).

### Inhibition of apoptotic pore formation by altering the lipid content of mitochondrial membranes

The specific release of lysophospholipids during pore formation raises the possibility that microdomains enriched for lysophospholipids contribute to pore formation. For example, lysophospholipids induce positive membrane curvature ^55^ and can limit the size and distribution of ordered lipid domains ^56^. These effects are even more pronounced if fatty acid chains are long (>20 residues) and polyunsaturated ^55, 56^. Cholesterol is an important regulator of membrane domains ^57^ and can counteract the positive curvature of lysophospholipids ^56^. As cholesterol can also inhibit BAX-mediated pore formation ^35, 36^ we next asked if addition of cholesterol can block BAK-mediated pore formation. We compared cholesterol with oxysterol 7-ketocholesterol (7KC) (Fig. S3c) ^58, 59^ that contains an additional ketone group which can lead to preferential expansion of ordered (cholesterol) versus disordered (7KC) domains ^57^. Both sterols can induce ordered (i.e. tightly packed) membrane domains when used at high concentrations ^57^.

Cholesterol and 7KC were incubated with mitochondria as water-soluble complexes with methyl-β-cyclodextrin (mβCD) (see Methods) ^58^. Pre-incubation of MLM with the vehicle only control (mβCD-EtOH) had no effect on cytochrome *c* release (Fig. 6a). In contrast, addition of mβCD-cholesterol or mβCD-7KC prevented cytochrome *c* release, with cholesterol appearing as a slightly more potent inhibitor than 7KC (Figs. 6a, b).

**Figure 6.**
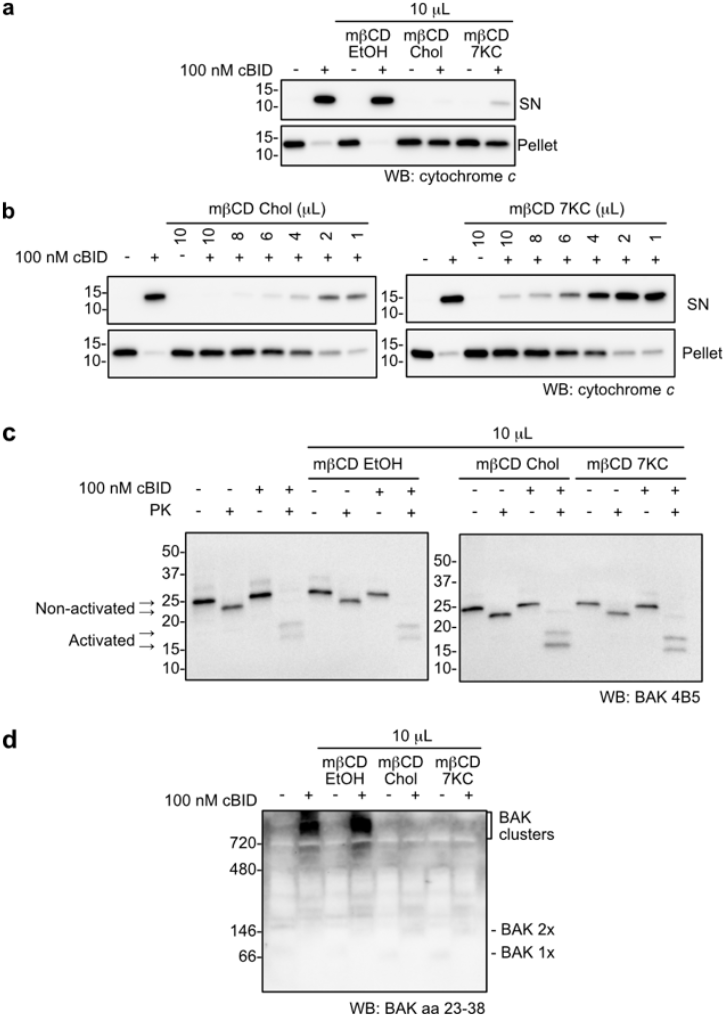
Cholesterol inhibits apoptotic pore formation in mouse liver mitochondria. WT MLM were incubated with the lipid carrier mβCD loaded with EtOH, cholesterol or 7KC, prior to incubation with cBID (100 nM) at 37°C for 30 min, and fractions analysed, as in Fig. 1. **(a)** Cytochrome *c* release was blocked by pre-incubation with cholesterol or 7KC. **(b)** Cholesterol and 7KC exhibit similar concentration-dependent inhibition of cytochrome *c* release. **(c)** BAK activation by cBID was not inhibited by cholesterol, as shown by susceptibility to proteinase K. Aliquots removed prior to fractionation were treated with proteinase K and immunoblotted with antibody to the BAK BH3 domain (clone 4B5). **(d)** BAK dimers failed to assemble into higher order BAK clusters in the presence of cholesterol or 7KC. The pelleted mitochondrial fractions from (a) were run on Blue Native PAGE and immunoblotted for BAK (aa23-38). Data are representative of three (a, d) or two (b, c) independent experiments.

To interrogate if cholesterol might be acting directly on BAK *en route* to pore formation, the impact of cholesterol and 7KC on BAK activation and BAK clustering was assessed. BAK was still activated by cBID, as shown by increased susceptibility to proteolysis by proteinase K (Fig. 6c) and loss of the monomeric nonactivated BAK (1X band; Fig. 6d). Notably, high molecular weight BAK clusters were absent (Fig. 6d) despite evidence of BAK homodimers migrating at 146 kD (Fig. 6d). Thus, cholesterol and 7KC do not interfere with BAK activation or dimer formation, but they do stop homodimers aggregating into high molecular weight complexes.

The ability of cholesterol to prevent the clustering of BAK homodimers (Fig. 6c) may explain its ability to prevent cytochrome *c* release (Figs. 6a,b). Due to its wide-ranging effects on membranes, cholesterol may also act downstream e.g. by inhibiting lipid microdomains and bilayer deformation (see Discussion). Nevertheless, the strong inhibition by cholesterol argues that the interplay between proteins and lipids is crucial to apoptotic pore formation in native mitochondria.

## DISCUSSION

To examine both the protein and lipid changes that drive apoptotic pore formation we used freshly isolated mitochondria as they contain endogenous levels of BAK residing in a native membrane environment. Upon apoptotic pore formation BAK remained in the mitochondrial pellet, while a specific group of lipids, namely long chain polyunsaturated lysophospholipids (LPLs), were released. To our knowledge, this is the first report of membrane phospholipids being released during apoptotic pore formation. The specificity of the lipid signature suggests sorting of these lipids into microdomains to drive a localised positive membrane curvature and subsequent lipid shedding (possibly as micelles) in concert with membrane rupture. Furthermore, a causal role for lipid microdomains in pore formation was suggested by the ability of cholesterol to block cytochrome *c* release and BAK clustering.

The lipid particles released with pore formation appear distinct from other recorded types of mitochondrial lipid particles such as structures positive for outer membrane (SPOTs)^60^, mitochondrial-derived vesicles (MDVs) ^61, 62^ and yeast mitochondrial-derived compartments (MDCs) ^63-66^ that are involved in quality control in response to various stresses. Those particles contain a range of proteins, and MDVs contained a lipid profile similar to the outer membrane ^61^. Note that the isolated mouse liver mitochondria prepared herein have undergone fission, which could preclude the release of large lipid particles such as those observed in cells.

The methodology adopted for lipidomics was designed to detect low levels of lipids released from mitochondria. The method involved initial sedimentation of the mitochondria by gentle centrifugation (10,000 *g*), and recovery of released material in the supernatant (SN^10^) by a second ultracentrifugation step (100,000 *g*). Lipidomics of this secondary pellet fraction (Pellet^100^) allowed removal of buffer constituents (e.g. HEPES) that interfere with ion detection during mass spectrometry, and also concentrated the SN^10^ samples. While sedimentation at 100,000 *g* may not capture the smallest and least dense lipid particles, good coverage of the lipids in our targeted panel was achieved as a large proportion of lipids in the targeted lipidomics panel (348/495 lipids) were detected in Pellet^100^. Importantly, comparison of apoptotic and non-apoptotic samples identified very specific lipid shedding events in response to apoptosis. Most striking was the over-representation of long chain polyunsaturated lysophospholipids (LPLs) upon BAK-driven pore formation. This lipid signature is consistent with BAK activation causing formation of distinctive lipid microdomains, which are unstable and vulnerable to fissure.

As a counterpoint to the LPL release driven by BAK activation, PLA2 treatment released lysophospholipids of all chain lengths and all levels of saturation. The PLA2 family of enzymes act at the face of a membrane to cleave di-acyl phospholipids at the sn-2 acyl linkage generating free fatty acids and lysophospholipids (Fig. S3a and reviewed ^49-51^). Interestingly, PLA2 promoted microdomain formation and vesiculation in GUVs ^67^ and degradation of supported lipid bilayers ^68^. Given the increased representation of lysophospholipids in Pellet^100^ following both PLA2 treatment and BAK-mediated pore formation, there may be mechanistic overlap in lipid release by the two treatments. However, a key distinction is that PLA2 enzymatic activity increases lysophospholipids, whereas BAK activation appears to drive the redistribution of existing lysophospholipids into microdomains.

Lysophospholipids are non-lamellar lipids due to their cone-shaped architecture (Fig. S3a). This shape confers a preference for adopting positive membrane curvature ^55^, for localising at the edge of toroidal pores ^69^ and for forming micelles in solution ^70^. In coarse grain simulations asymmetric incorporation of lysophosphatidylcholine induced significant membrane curvature ^71^. Moreover, simulations show cone-shaped lipids coalesce into domains that burst the membrane leaving behind a single toroidal pore lined by cone-shaped lipids ^72^. Thus, release of lysophospholipids that are long chain and polyunsaturated during apoptosis suggests these lipids become enriched within dedicated microdomains where positive curvature can cause lipid shedding and pore formation (Fig. 7, Movie S1).

**Figure 7.**
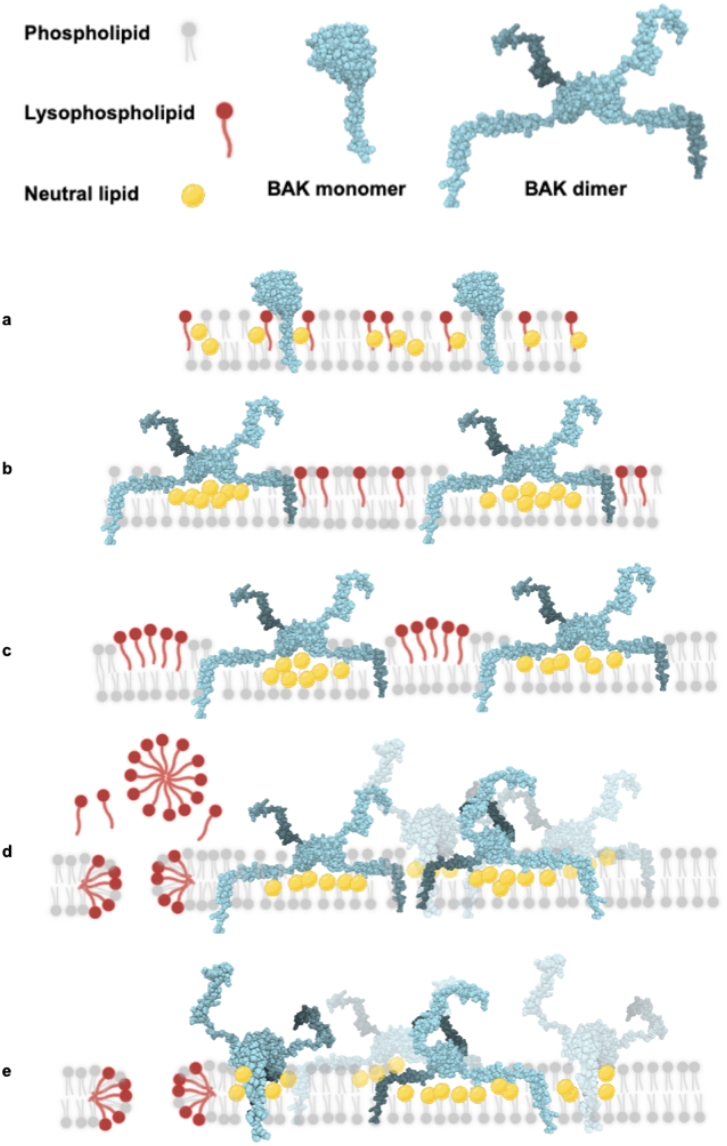
BAK-lipid microdomain model of apoptotic pore formation. **a)** In healthy mitochondria, BAK monomers are distributed on the MOM surface surrounded by a mixture of lipids (phosphoplipids, grey). Long chain polyunsaturated lysophospholipids (LPLs, red, <1% total) are distributed in the membrane and associated with neutral lipids (e.g. TAGs and ceramides, yellow). **b)** After apoptotic signalling, BAK dimers form and sink into the outer leaflet, displacing lipids with large polar headgroups (e.g. phospholipids) and recruiting neutral lipids under each BAK dimer. Any associated lysophospholipids also become more mobile. **c)** Separate microdomains of LPLs and BAK dimers form. LPL microdomains drive positive curvature and initial delamination of the two leaflets. **d)** Growth of microdomains driven by membrane stress. Continued LPL aggregation drives severe positive membrane curvature, with non-bilayer LPLs liberated from the membrane (possibly as a micelle), and leaving behind a lipidic pore partially lined with LPLs. **e)** Pore stabilization and enlargement. Free movement of lipids between leaflets can relieve membrane tension. In addition, BAK dimers remain anchored in the membrane via their transmembrane domains, with the less restrained core region dragged adjacent to the pore edge. Created with BioRender.com

Lipidomics also revealed a subtle depletion of certain neutral lipids (TAGs and ceramides) in the Pellet^100^ samples following apoptotic signaling. Neutral lipids like TAGs and ceramides lack polar headgroups (Fig. S3b) and thus reside more centrally in the hydrophobic interleaflet space of a membrane bilayer. For instance, TAGs readily form non-lamellar “lens structures” within membrane bilayers ^73^. The depletion of these neutral lipids in our Pellet^100^ samples suggests their retention in mitochondria during pore formation. Retention of TAGs is consistent with recent molecular dynamics simulations of TAGs aggregating beneath the BAK α2-α5 core dimer in response to membrane deformation by the dimer ^18^. Although we have no direct measure of TAG association with BAK clusters in our system, recruitment of TAGs under BAK dimers provides a plausible mechanism for BAK dimer-driven lipid redistribution and aligns with retention of BAK in the mitochondrial fraction (Fig. 2).

Interestingly, another feature of certain neutral lipids is their capacity to co-segregate with lysophospholipids by packing closely with the lysophospholipid acyl chains underneath the lysophospholipid headgroup ^74^. Presuming that neutral lipids (e.g. TAGs and ceramides) tend to co-segregate with lysophospholipids in healthy mitochondria, any concentration of neutral lipids under BAK dimers may leave the lysophospholipids “orphaned”, prompting formation of lysophospholipid microdomains with positive curvature. Together, the effects of neutral lipids segregating with BAK dimers and of lysophospholipids forming new positively curved microdomains could have a dramatic impact on the integrity of the membrane bilayer.

To test the impact of altered lipid composition on pore formation, mitochondria were pre-incubated with exogenous cholesterol. Mitochondrial membranes maintain relatively low levels of cholesterol ^42, 45, 48^, with increased levels linked to various pathologies (reviewed in ^75^). Cholesterol has many effects on model membranes, including the ordering of phospholipid acyl chains (formation of “lipid rafts”), lipid phase separation and increased bilayer thickness (reviewed in ^76, 77^). Here we show that in MLM, added sterols blocked BAK dimer clustering and MOMP (Fig. 6). Given the pleiotropic roles of cholesterol in membrane behaviour, cholesterol treatment may be inhibiting various stages of BAK-lipid mediated pore formation.

We first considered if BAK activation and dimerization could be inhibited by cholesterol. Previous studies with BAX in liposomes and isolated mitochondria showed that added cholesterol increased lipid order and inhibited early steps in BAX function including translocation to membranes, membrane insertion and activation (conformation change) ^35, 36. 75^. We found no such inhibition of BAK activation or dimer formation by cholesterol, and this aligns with BAK being constitutively present in the MOM as a tail-anchored protein ^7^, in contrast to BAX ^8-10^.

Next, we considered how cholesterol might block BAK dimer clustering. Formation of oligomers or multimers of BAK (or BAX) dimers has long been considered important for pore formation, yet there is no well-defined protein:protein interface between dimer subunits that explains oligomerisation. Several lines of evidence suggest that the membrane bilayer itself promotes clustering of BAK dimers ^15, 19, 26^, supported here by the ability of cholesterol to inhibit clustering. According to modelling studies, clustering of membrane proteins may result when the membrane bilayer attempts to minimise hydrophobic mismatch between the protein and surrounding lipids ^78^. Thus, following shallow insertion of the BAK core dimer, cholesterol may minimise mismatch by being recruited to the dimers (as per TAGs) or by increasing the membrane thickness surrounding each dimer. Cholesterol’s ability to create ordered domains with reduced lateral mobility in the bilayer may also inhibit protein clustering. A complementary phenomenon is that increasing phospholipid unsaturation increases membrane fluidity ^79^ and promotes MOMP ^39^.

Downstream of BAK clustering, cholesterol may directly inhibit lysophospholipid microdomains and/or their collapse into lipidic toroidal pores. For example, increased cholesterol prevented curvature stress induced by lysophospholipids in simulations and AFM studies ^56^. Cholesterol and other sterols can co-segregate with lysophospholipids via hydrophobic interactions ^80, 81^. Thus, especially considering that cholesterol levels in mitochondria are relatively low ^42, 45, 48^, a small increase may counter effects of other neutral lipids (e.g. TAGs, ceramides). In summary, there are several modes by which cholesterol treatment could hinder pore formation in our model system. Future simulations of membranes containing a wide repertoire of lipids (e.g. lysophospholipids that are saturated or unsaturated, TAGs, ceramides and cholesterol) may help delineate which mode(s) of action accounts for the robust inhibition of pore formation by cholesterol.

Identifying the topology of BAK dimers in relation to the membrane bilayer, to other dimers, and to a toroidal pore, has been of significant interest in terms of understanding the mechanism of pore formation and how to regulate apoptotic cell death. A range of studies indicate that a large part of each BAK dimer interacts with the membrane, including transmembrane insertion of the C-terminal α9-helices at either end of the dimer, and outer leaflet insertion by the amphipathic α2-α5 core and α6-helices (reviewed in ^25^, Fig. 7, Movie S1). This dimer topology is consistent with retention of BAK in the mitochondrial fraction rather than release with lysophospholipids. Thus, BAK retention in the bulk pellet indicates that apoptotic pore formation is distinct from the “carpet model” of membrane disruption reported for certain types of pore-forming peptides ^82^, and also excludes a model like the “cookie cutter” pores formed by the lytic cell death regulator NINJ1 in which the protein subunits and the membrane disc enclosed within that ring are both liberated from the membrane ^83^.

The topology of each BAK dimer in relation to other dimers, and to the pore itself, is less clear. Microscopy suggests that within each apoptotic mitochondrion most of BAK forms oligomers (i.e. clusters), but only a portion positions at the pore edge ^26, 84, 85^. This implies that clustering precedes pore formation, but also implies that biochemical assays will be monitoring a mixture of two (or more) populations of BAK dimers. Distinct topologies may exist for BAK dimers in clusters and at the pore edge based on flexibility in the latch regions (e.g. α5-α6 and α8-α9 linker regions). Dimers clustered on the membrane surface may adopt the in-plane orientation ^21^, followed by re-positioning at the rim of a newly formed pore. Further re-positioning of the α2-α5 core across pore edge, as in the clamp model, would involve one α9 anchor becoming inverted across the bilayer ^86, 87^.

Our findings lead us to a BAK-lipid microdomain model of apoptotic pore formation (Fig. 7, Movie S1). As BAK homodimers embed in the membrane surface, they drive membrane thinning and lipid redistribution, causing lipids without polar headgroups (neutral lipids such as TAGs and perhaps ceramides) to focus beneath the dimer. The “orphaned” LPLs then coalesce into a microdomain minimising hydrophobic mismatch at the interface of individual LPLs and the surrounding bilayer. In parallel, BAK dimers coalesce into microdomains (clusters) to minimise hydrophobic mismatch at the interface of individual BAK dimers and the surrounding bilayer. When a critical density of LPLs is attained, the localised positive curvature results in membrane rupture and release of LPLs (possibly as a micelle), leaving behind a small unstable lipid-lined toroidal pore. The initial pore may be stabilised and enlarged (or distorted to create a tear) by two events: flow of lipids from the outer to the inner leaflet around the pore edge to neutralise membrane tension; and BAK homodimers positioning at the pore rim.

This BAK-lipid microdomain model is consistent with other findings from native mitochondrial membranes including BAK multimers adopting a variety of architectures (i.e. arcs, lines, rings, clusters) and an essential protein:protein contact between dimers has not been shown. The model highlights the requirement for lipid reorganisation to effect BAK-mediated pore formation. During BAX-mediated pore formation also, a temperature-dependent “lag phase” evident in outer membrane vesicles obtained from mitochondria may relate to lipid microdomains forming prior to membrane rupture ^32^.

In conclusion, examining endogenous BAK protein in its native mitochondrial environment has enabled a broad survey of lipids liberated from the membrane upon formation of the apoptotic pore. We have identified for the first time a lipid signature of long chain polyunsaturated lysophospholipids released during apoptotic pore formation. Data showing that exogenous lipids can prevent BAK dimer clustering highlights the important interplay of BAK and lipids *en route* to pore formation. Collectively, our data support the lipidic nature of BAK-dependent pores, with the BAK-lipid microdomain model providing a possible mechanism for membrane forces driving both BAK clustering and pore formation. Future testing for lysophospholipid release from toroidal pores formed by other proteins, and from other native and model membranes, may reveal if the lipid signature we have observed for apoptotic pores in mouse liver mitochondria is a common feature of toroidal pore formation.

## METHODS

### Isolation of mouse liver mitochondria

Mouse liver mitochondria (MLM) were isolated from male wildtype *C57BL/6* or *Bak*^-/-^*C57BL/6* mice ^2^ (ranging 74 to 155 days old, wildtype n=6, *Bak*^-/-^ n=6) that were bred and maintained at the Walter and Eliza Hall Institute of Medical Research Animal Facility. All animal experiments were approved by the WEHI Animal Ethics Committee (WEHI internal ethics number 2022.014) and were conducted in accordance with the Prevention of Cruelty to Animals Act (1986) and the Australian National Health and Medical Research Council Code of Practice for the Care and the Use of Animals for Scientific Purposes (1997). Trained Bioservices staff performed euthanasia of adult mice by cervical dislocation. Mouse livers were collected in chilled sucrose solution (300 mM sucrose, 10 mM Tris HCl, pH7.4, 1 mM EDTA). Isolated mitochondria were prepared from livers as described previously ^46^. The concentration of MLM was estimated by absorbance at 280 nm.

### Reagents

Recombinant caspase-8 cleaved human BID (cBID) was prepared as per ^46^ and aliquots stored at -80°C before thawing on ice. Phospholipase A2 (PLA2) from bee venom (Sigma Aldrich) was prepared as a stock of 1mg/mL and stored at -20 °C. Methyl β cyclodextrin (mβCD) (Sigma Aldrich) is a lipid carrier commonly used at high concentrations to extract cholesterol, or at lower concentrations (as per this study) to solubilise sterols for delivery to membranes ^58^. Methyl β cyclodextrin (mβCD) was loaded with cholesterol (Sigma Aldrich) or 7-ketocholesterol (7KC) (Sigma Aldrich) as described ^58^. Briefly, 5% w/v mβCD (50 mg/mL, 400 μL aliquots) was pre-warmed to 80°C on a heatblock for 5 min. Sterols dissolved in ethanol at 15 mg/mL were added to the heated mβCD, 10 μL added at 5 min intervals mixing with vortex with each addition and returned to heat, and repeated four times for a total of 40 μL sterol added per 400 uL preparation to yield 1.5 mg/mL of each sterol in 50 mg/mL mβCD. To generate a “vehicle-only” control, an ethanol only loading of mβCD was also prepared (mβCD-EtOH) in parallel. Sterol reagents were stored at 4°C protected from light.

### MLM incubations

Freshly prepared MLM (used within 3 hours of isolation) were kept on ice before dilution in mitochondrial assay buffer containing 100 mM KCl, 2.5 mM MgCl2, 100 mM sucrose, 20 mM HEPES/KOH at pH 7.5, protease inhibitor cocktail (Roche) and 8 μg/mL pepstatin A (Sigma Aldrich). For lipidomic studies, samples were incubated for 45 min at 37°C with either 100 nM cBID or 5 μg/mL or 50 μg/mL PLA2. For sterol treatments, 200 μL aliquots were pre-incubated with 1-10 μL 1.5 mg/mL mβCD-cholesterol, mβCD-7KC or mβCD-EtOH at 37°C for 30 min, then treated with 100 nM cBID for 30 min. For EDTA inhibition assays, samples were pre-incubated with 20 mM EDTA at 37°C for 30 min, prior to addition of cBID or PLA2.

### MLM cytochrome c release assay

To monitor mitochondrial permeabilisation, pellet and supernatant fractions from a 10,000 *g* 5 min fractionation were analysed by SDS PAGE. Fractions were resuspended reducing 2x SDS sample loading buffer (150 mM Tris pH 6.8, 1.2% (w/v) SDS, 30% (v/v) glycerol, 0.018 mg/mL bromophenol blue, 5% v/v 2-mercaptoethanol). Samples were boiled for 3 min, then resolved by SDS PAGE (12% TGX gels, BioRad) and transfered to 0.2 μm nitrocellulose membrane (Bio-RAD) via wet transfer (25 mM Tris, 192 mM Glycine, 20% v/v Methanol). Membranes were blocked in 5% w/v skim milk in TBS with 0.1% v/v Tween and immunoblotted for cytochrome *c* (mouse monoclonal antibody, clone 7H8.2C12, #556433, BD Biosciences) or HSP60 (rabbit polyclonal antibody, #A302845A, Thermo Fisher Scientific).

### MLM BAK activation assay with proteinase K proteolysis

BAK activation status was monitored via susceptibility of the BAK protein to digestion with proteinase K (PK) as per ^88^. Whole mitochondrial fractions were pre-chilled on ice for 15 min, prior to addition of proteinase K (30 μg/mL final) and incubation for 20 min on ice. The reaction was stopped with the addition of PMSF (2 mM final), then an equal volume of reducing 2x SDS sample loading buffer added to each sample. Samples were resolved by SDS PAGE (12% TGX gels, BioRad) and transfered to 0.2 micron nitrocellulose via wet transfer (25 mM Tris, 192 mM Glycine, 20% v/v Methanol). Membranes were blocked in 5% w/v skim milk in TBS with 0.1% v/v Tween and immunoblotted for BAK 4B5 (rat monoclonal antibody, clone 4B5, WEHI mAb facility).

### Blue Native PAGE

Higher order complexes of BAK protein were measured using Blue Native PAGE. Samples were analysed by Blue Native PAGE (BNP) to preserve the native interface within BAK dimers and allow assessment of clustering of multiple BAK dimers. Membrane fractions were isolated by centrifugation at 10,000 *g* for 5 min. Supernatants were discarded and membrane pellets were solubilised in 20 mM Bis-Tris pH 7.4, 50 mM NaCl, 10% v/v glycerol, 1% w/v digitonin, 10 mM DTT and incubated on ice for 1 hr. A second centrifugation of 16,000 *g* for 5 min was used to sediment any insoluble material. Native Sample buffer (Life Technologies) and Coomassie Additive (Life Technologies) were added to the soluble supernatants. In some instances, detection of BAK dimers was enhanced by supplementation with 5 mM EDTA. Samples were resolved with Novex 4–16% Native PAGE 1.0 mm 10 well gels (Life Technologies). Western blot transfer to 0.2 μm PVDF membrane (Thermo Fisher Scientific) was performed via wet transfer (25 mM Tris, 192 mM Glycine, 20% v/v Methanol, supplemented with 0.037% w/v SDS). PVDF membranes were de-stained with 10% v/v acetic acid 30% v/v ethanol, then further de-stained in methanol and rinsed thoroughly with dH2O before immunoblotting. Membranes were blocked in 5% w/v skim milk in TBS with 0.1% v/v Tween, and immunoblotted for BAK aa23-38 (clone DF9) (rabbit polyclonal antibody, B5897, Sigma-Aldrich, Castle Hill, NSW, Australia).

### Western blot secondary antibody detection and image capture

For secondary detection, membranes were incubated with Horseradish peroxidase conjugated IgG secondary antibodies; anti-rabbit (4010–05, Southern Biotech), anti-rat (3010–05, Southern Biotech) and anti-mouse (1010–05, Southern Biotech). Immobilised horseradish peroxidase was detected with Immobilon Forte Western HRP substrate (WBLUF0500, Millipore, Billerica, MA, USA), images captured with the ChemiDoc MP System (Bio-RAD, Hercules, CA, USA) and signal intensity measured with Image Lab 6.1 software (Bio-RAD).

### MLM fractionation for lipidomic analysis

A small volume of each sample for lipidomic analysis (50 μL from 1.5 mL total) was prepared by centrifugation at 10,000g for 5 min, and pellet and SN fractions collected. Fractions were resolved by SDS-PAGE and cytochrome *c* release measured (as described above). The remainder each treatment sample (∼1.45 mL) was subjected to a 2-step fractionation procedure. Samples were first separated by centrifugation at 10,000 *g* (10,000 rpm) 5 min at 4°C, and the supernatant (SN^10^) retrieved. The SN^10^ fraction was then applied to pre-weighed tubes (Beckman Coulter, maximum volume 1.5 mL #357448). Samples were then separated by a centrifugation at 40,000 rpm (TLA-55 angle rotor S/N 19U1568 in Beckman Coulter Optima Max XP Ultra Centrifuge, r^max^ 98,600*g*) for 20 min at 4 °C. Supernatants were removed and pellets weighed and placed on wet ice. The mass of samples ranged from 18.5 mg to 33.9 mg from pooled duplicate pellets (Supp. Table 1). Pellets were stored at -80°C prior to being pooled during extraction and analysis by Metabolomics Australia (Bio21 Institute, Melbourne, VIC, Australia).

### Lipidomic profiling

Targeted lipidomic profiling was conducted for samples fractionated from isolated liver mitochondria from age-matched adult male mice (wildtype C57BL/6 n = 6, *Bak* ^-/-^ n = 6, male, ages ranged from 74 to 156 days old, 3 treatment groups per mouse with a total of 36 samples processed for lipidomics (Supp. Table 1). Internal standards comprising a mixture of PC(19:0/19:0), PE-D31, PG (17:0/17:0), TG-D5 (Avanti Polar Lipids, Alabama, USA) were added (10 mg/L) to the stock extraction solution. The extraction solution containing internal standards were used for all sample extractions to assess sample processing and instrument performance. Ice cold 1:9 chloroform:methanol (600 μL) and internal standards were added to sample pellets. Suspensions were transferred to 2 mL microcentrifuge tubes, then 1 mL 100% (v/v) chloroform added to obtain a final ratio of 2:1 chloroform:methanol. Samples were vortexed for 30 sec, then mixed at 950 rpm for 5 min at 10 °C in an Eppendorf Thermomixer (Eppendorf South Pacific Pty Ltd., Macquarie Park, Australia). Samples were centrifuged at 15,000 rpm for 5 min at 10 °C and supernatants collected. Supernatants were completely dried in a vacuum concentrator to complete dryness (100 μL for 15 cycles, Christ® RVC 2–33, Martin Christ Gefriertrocknungsanlagen GmbH, Osterode am Harz, Germany). Dried samples were reconstituted in methanol:water-saturated butanol (100 uL, 1:9, v/v). To assess instrument performance, pooled biological quality control samples (PBQCs) were prepared by combining 10 μL aliquots of each sample extract. Samples were injected (2 μL volume) in a random order for analysis by LC-QQQ-MS. PBQCs were injected at intervals, after every 4 samples.

Extracted lipids were processed and detected by Metabolomics Australia (Bio21 Institute, Melbourne, VIC, Australia) as previously described ^89, 90^ with an Agilent 1290 liquid chromatography (LC) system and Triple Quadrapople 6490 mass spectrometer (MS, Agilent Technologies Australia, Mulgrave, Australia).

A curated list of 348 metabolites (from the targeted 495 metabolites in the Mass Hunter Database/Mass Hunter Quantitative Analysis) was obtained and data imported into MetaboAnalyst 5.0 for initial quality control checks and analysis by Metabolomics Australia (Bio21 Institute, Melbourne, VIC, Australia). This curated dataset was subject to further downstream statistical analysis as follows.

### Statistical analysis of lipidomic data

Relative abundance of lipids were analysed in *R* ^91^ using *limma* ^*92*^,*Glimma* ^*93*^ and *edgeR* ^94^. After removing pooled biological quality control (PBQC) samples, quantile normalization ^95^ of log2-counts was performed. Linear models ^96^ with sample-specific weights ^97^ were fitted to summarise the data from each experimental group (WT Untreated, WT cBID, WT PLA2, KO Untreated, KO cBID and KO PLA2) while accounting for variation between sample collection dates. Pairwise contrasts between the experimental groups were estimated, and differential lipid abundance was assessed using moderated *t*-statistics ^96^. Lipids were ranked according to their false discovery rate (FDR) ^98^, and those with a FDR < 0.05 were considered differentially abundant. Figures were plotted with functions in the *limma, ggplot2* ^99^ and *ggrepel* ^100^ packages.

### Lipid set enrichment analysis

Lipids were classified according to headgroup and enrichment of classes with at least 3 lipids was tested using ROAST ^52^. LION/web analysis was performed in “target-list mode”. The top 26 lipids (raw *p*-value < 0.2) from the WT cBID vs WT Untreated contrast with positive log2fold-change were input as the “lipid target list” and the full list of lipids detected were input as the “lipid background list”.

## DATA AVAILABILITY STATEMENT

Lipidomic data, analysis code and interactive summary plots are available via GitHub at https://github.com/mritchie/BAKLipidomicAnalysis/.

## ANIMAL ETHICS APPROVAL

All animal experiments were approved by the WEHI Animal Ethics Committee (WEHI internal ethics number 2022.014) and were conducted in accordance with the Prevention of Cruelty to Animals Act (1986) and the Australian National Health and Medical Research Council Code of Practice for the Care and the Use of Animals for Scientific Purposes (1997).

## ACKNOWLEDGEMENTS

We thank Professor Brian Smith for helpful discussions.

## AUTHOR CONTRIBUTIONS

RTU: Conceptualization, Formal analysis, Supervision, Investigation, Visualization, Methodology, Writing—original draft, Writing—review and editing.

MER: (Lipidomics) Data curation, Software, Formal analysis, Visualization, Methodology, Writing—review and editing.

AWW, JPL: Investigation, Writing—review and editing. EU: Visualization, Writing—review and editing.

VKN, DPDS: (Lipidomics) Investigation, Methodology, Writing—review and editing. DS: Writing—review and editing.

RMK: Conceptualization, Supervision, Visualization, Methodology, Writing—original draft, Writing—review and editing.

## SUPPLEMENTARY MATERIAL

Supplementary materials are available online.

Supplementary Tables S1-S8

Supplementary Movie S1

Supplementary Figures S1-S4

