## Supplementary Materials for "A lipid signature of BAK-driven apoptotic pore formation"

### Supplementary Material

#### Table S1. Sample collection details.

**Table S2. Targeted lipidomics panel categorised by class.** The number and percentage of each lipid class detected in Pellet$^{100}$ fractions is also shown.

| Abbreviation | Lipid class | # Lipids in panel | # Lipids detected | % Lipids detected |
|---|---|---|---|---|
| PC(19:0/19:0)(IS) | PC internal standard | 1 | 1 | 100 |
| PE-D31(IS) | PE internal standard | 1 | 1 | 100 |
| PG 17:0 17:0 (IS) | PG internal standard | 1 | 1 | 100 |
| TG-D5 (IS) | TG internal standard | 1 | 1 | 100 |
| Acylcarnitine | Acylcarnitine | 12 | 3 | 25 |
| CE | Cholesterol esters | 27 | 13 | 48 |
| Cer | Ceramides | 41 | 18 | 44 |
| COH | Cholesterol | 1 | 1 | 100 |
| Des | Desmosterol | 6 | 0 | 0 |
| DG | Diacylglycerides | 20 | 19 | 95 |
| dhCer | Dihydroxyl ceramides | 6 | 3 | 50 |
| GM | Gangliosides | 8 | 0 | 0 |
| Hex1/2 Cer | Hexosyl Ceramides | 30 | 9 | 30 |
| LPC | Lysophosphatidylcholine | 27 | 26 | 96 |
| LPC-O | Ether linked Lysophosphatidylcholine | 10 | 7 | 70 |
| LPC-P | Plasminogen linked Lysophosphatidylcholine | 5 | 4 | 80 |
| LPE | Lysophosphatidylethanolamine | 7 | 7 | 100 |
| LPE-P | Plasminogen linked Lysophosphatidylethanolamine | 4 | 4 | 100 |
| LPI | Lysophosphatidylinositol | 4 | 4 | 100 |
| OH cholesterol | OH cholesterol | 4 | 0 | 0 |
| oxCE | Oxysterols | 2 | 0 | 0 |
| PC | Phosphatidylcholine | 45 | 39 | 87 |
| PC-O | Ether linked Phosphatidylcholine | 19 | 15 | 79 |
| PC-P | Plasminogen linked Phosphatidylcholine | 21 | 15 | 71 |
| PE | Phosphatidylethanolamine | 23 | 22 | 96 |
| PE-O | Ether linked Phosphatidylethanolamine | 11 | 9 | 82 |
| PE-P | Plasminogen linked Phosphatidylethanolamine | 38 | 25 | 66 |
| PG | Phosphatidylglycerol | 4 | 4 | 100 |
| PI | Phosphatidylinositol | 21 | 18 | 86 |
| PS | Phosphatidylserine | 7 | 6 | 86 |
| SM | Sphingomyelins | 34 | 25 | 74 |
| Sulfatide | Sulfatides | 6 | 0 | 0 |
| TG | Triacylglycerides | 44 | 44 | 100 |
| TG-O | Ether linked Triacylglycerides | 3 | 3 | 100 |
| Ubiquinone | Ubiquinone | 1 | 1 | 100 |
| Total | | 495 | 348 | 70 |

**Table S3. Relative yield of each lipid class from lipidomic assessment of WT untreated samples, compared to Ardail et al [45].**

| **Abbreviation** | **Lipid class** | **Ardail et al. outer membrane Lipid class as % total lipids** (means of 5 separate preparations in percent by weight of total lipids) | **Lipidomic raw values Lipid class as % total lipids** (means of 6 separate preparations in percent by counts of total lipids) |
|---|---|---|---|
| Acylcarnitine | Acylcarnitine | - | 0.002 |
| CE | Cholesterol esters | - | 0.187 |
| Cer | Ceramides | - | 0.808 |
| COH | Cholesterol | 7.1 | 0.006 |
| DG | Diacylglycerides | 0.3 | 0.331 |
| dhCer | Dihydroxyl ceramides | - | 0.003 |
| Hex1/2 Cer | Hexosyl Ceramides | - | 0.035 |
| LPC | Lysophosphatidylcholine | 0.4 | 1.219 |
| LPE | Lysophosphatidylethanolamine | 0.1 | 0.317 |
| LPI | Lysophosphatidylinositol | - | 0.005 |
| PC | Phosphatidylcholine | 40.9 | 66.136 |
| PE | Phosphatidylethanolamine | 26.8 | 12.557 |
| PG | Phosphatidylglycerol | - | 0.042 |
| PI | Phosphatidylinositol | 9.1 | 4.618 |
| **PS** | Phosphatidylserine | <0.1 | 0.939 |
| **SM** | Sphingomyelins | 1.8 | 6.487 |
| **TG** | Triacylglycerides | 0.7 | 6.307 |
| **Ubiquinone** | Ubiquinone | - | 0.002 |
| **Fatty acids** | Fatty acids | 8.2 | - |
| **Cardiolipin** | Cardiolipin | 4 | - |
| Total | | 99.4 | 100 |
| | "-"denotes lipids not assayed | | |

**Table S4. Lipid abundance represented as raw lipidomic output without normalisation.**

**Table S5. Lipid abundance represented as normalised lipidomic output.** Excludes pooled biological quality control samples and samples which did not pass cytochrome c release quality control.

**Table S6. Fold-change lipidomics analysis of each contrast.** Values are ranked by $\log_2$fold-change.

**Table S7. ROAST lipid set enrichment analysis.**

**Table S8. LION/web lipid set enrichment analysis.**

**Movie S1. Animation of the BAK-lipid microdomain model of apoptotic pore formation.**

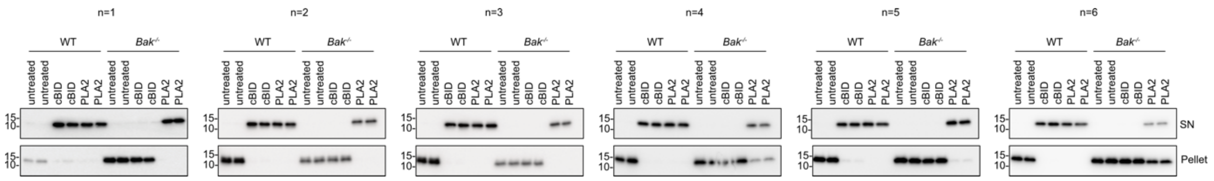


**Figure S1. Screening MLM incubations for cytochrome *c* release prior to processing for lipidomics.**

MLM were prepared in six separate experiments, each involving one WT and one *Bak*$^{-/-}$ mouse. Duplicate aliquots of MLM were prepared for each indicated treatment (untreated, 100 nM cBID or 5 $\mu$g/mL PLA2). After incubation for 30 min at 37°C , small separate aliquots were taken from each larger incubation volume, and all samples centrifuged at 10,000 *g*. The small aliquots of supernatant (SN) and mitochondria (Pellet) were analysed by western blot for cytochrome *c*. The remaining large bulk volume of each duplicate SN sample was concentrated further into separate 100,000 *g* pellets (as illustrated in Fig. 2a) and stored at -80°C until duplicate pellets were pooled and then processed for lipidomic analysis. Incomplete cytochrome *c* release was evident in some PLA2 treated samples (*Bak*$^{-/-}$ #4 and *Bak*$^{-/-}$#6).

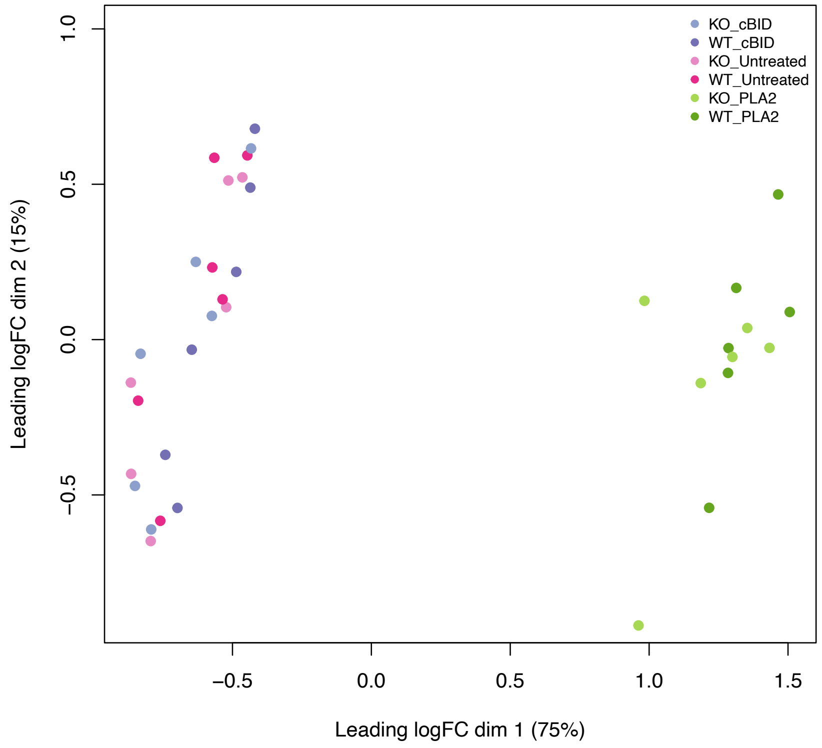


**Figure S2. PLA2 digested mouse liver mitochondria exhibit a distinct lipid profile.**

Multi dimensional scaling (MDS) plot of lipidomic output shows that most of the variation between samples (dimension 1) separates PLA2-treated (right-hand side) samples from the Untreated and cBID samples (left-hand side).

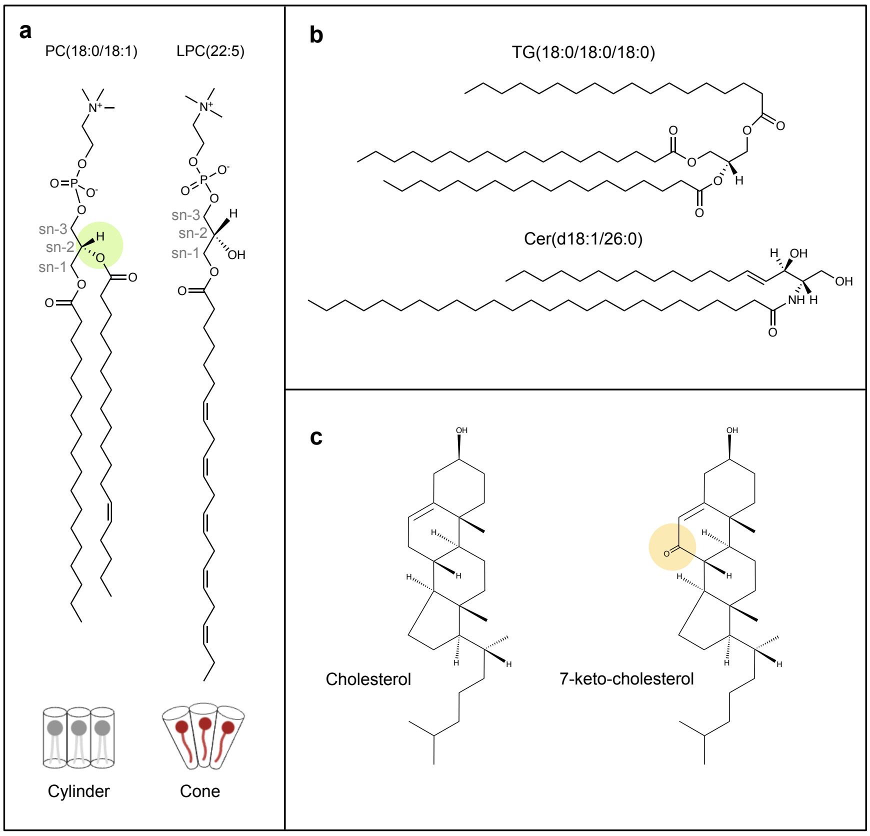


**Figure S3. Chemical structures of lipids.**

**(a)** Examples of phosphatidylcholine (PC) and lysophosphatidylcholine (LPC) molecules. Phospholipids include a phosphate headgroup and acyl (fatty acid) chains of variable length and saturation. Note that PLA2 specifically hydrolyses di-acyl phospholipids (e.g. PC) at sn-2 (green highlight) to yield a monoacyl lysophospholipid and a free fatty acid. The 3D geometry of di-acyl phospholipids is typically cylindrical, wheras mono-acyl phospholipids is conical. Structures of PC(18:0/18:1)(LMGP01010753) and LPC(22:5) (LMGP01050143) are shown.

**(b)** Two examples of neutral lipids**.** Lipids most frequently retained in the pellet with apoptotic pore formation were lipids without charged headgroups; TG (18:0_18:0_18:0)( LMGL03010002) and ceramide (d18:1_26:0)( LMSP02010011).

**(c)** Cholesterol (LMST01010001) and 7-ketocholesterol (7KC) (LMST01010049) differ by one ketone (yellow highlight), which may increase the hydrophilic contacts of 7KC relative to cholesterol and thus alter depth of insertion into a phospholipid bilayer and exhibit different preferences for ordered (cholesterol) verus disordered (7KC) membranes. However, at high concentrations, both sterols can induce the formation of ordered membrane domains [57].

Structure coordinates acquired from www.lipidmaps.org (IDs are indicated (LM#).

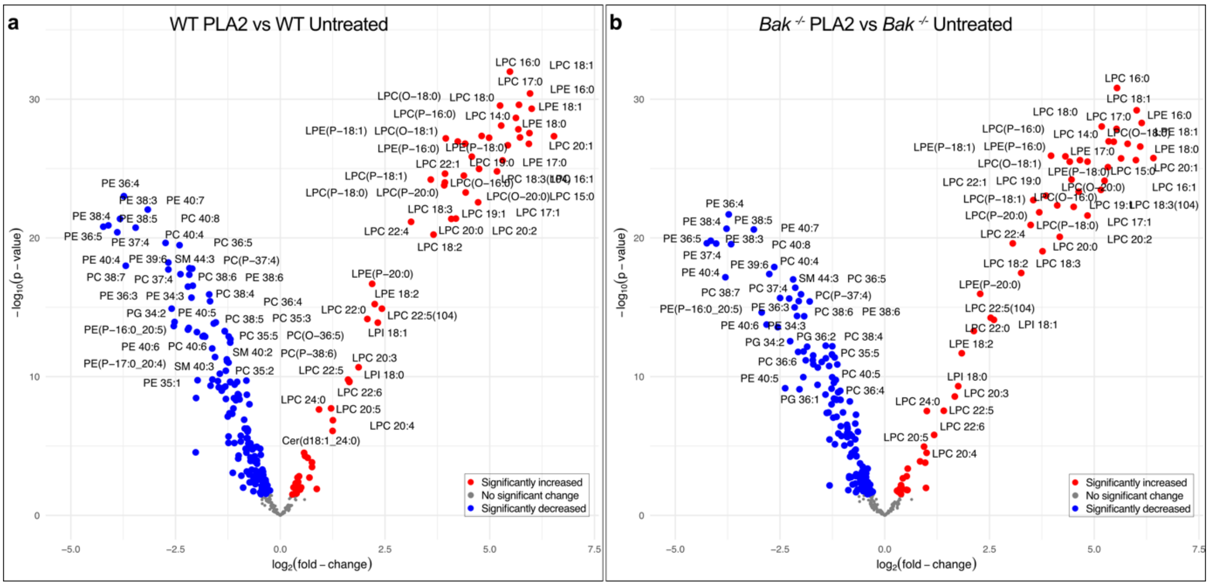


**Figure S4. PLA2 digested mouse liver mitochondria release lysophospholipids typical of PLA2 digestion.**

Volcano plots illustrate comparison between Pellet$^{100}$ fractions from **(a)** WT PLA2 versus WT Untreated samples and **(b)** *Bak$^{-/-}$* PLA2 versus *Bak$^{-/-}$* Untreated samples. Lipids with FDR <0.05 are highlighted in red (increased) or blue (decreased). Lipid $\log_2$(fold-change)(x-axis) is plotted against (-$\log_{10}$(p-value)(y-axis). A wide range of LPC and LPE lipid species are highly enriched in the supernatant of PLA2 treatments of both genotypes, and a range of potential substrate PC and PE species were depleted in parallel.
